# RERconverge Update: Runtime Reduction and Analysis Function Overhaul

**DOI:** 10.64898/2026.06.06.730612

**Authors:** Guillermo L. Hoffmann, Emily E. K. Kopania, Michael Tene, Amanda Kowalczyk, Ruby Redlich, Andreas R. Pfenning, Wynn K. Meyer, Maria Chikina, Nathan L. Clark

## Abstract

**Motivation:** Convergent evolution, or the independent acquisition of similar phenotypes in distinct lineages, provides a powerful framework for investigating genomic changes associated with a phenotype. This paper details an update to RERconverge, a powerful R package that tests for associations between gene relative evolutionary rates (RERs) and convergent phenotypes to infer genomic regions associated with traits or selective pressures. We introduce new customizable analysis choices and scalable and efficient algorithms that can process larger genomic datasets, a critical improvement as genomic data become available for more species.

**Results:** Modifications to core functions in the RERconverge pipeline resulted in an immense speedup (by a factor of up to 28.6). The function that tests for associations between phenotypes and RERs has been expanded to include two new analytical methods for outlier control; we also provide here a summary of the statistical tests users can perform, along with their use cases.

**Availability and implementation:** The code and walkthrough vignettes for the package are available at https://github.com/nclark-lab/RERconverge.

**Contact:** Nathan L. Clark; Maria Chikina

## Introduction

Repeatedly across the tree of life, distant species have acquired similar characteristics through convergent evolution. Some examples include the independent loss of flight in birds (Sackton et al. 2019), the repeated evolution of carnivorous plants (Ellison and Gotelli 2009), and the independent transition to aquatic life in three mammalian clades (Chikina, Robinson, and Clark 2016). These convergent phenotypes are indicative of similar selective pressures acting on independent lineages. In these cases, we often see similar genes or molecular pathways changing across these lineages in response to similar selective pressures. For example, cetaceans and bats independently evolved echolocation, and the auditory gene *Prestin* is under selection in both lineages (Li et al. 2010). Investigating convergent molecular changes associated with phenotypic adaptations and selective pressures has also helped uncover novel genes associated with traits such as eyesight (Partha et al. 2017) and longevity (Kowalczyk et al. 2020).

A powerful approach to study the molecular basis of convergent phenotypes is to examine the lineage-specific relative evolutionary rates (RERs) of genes or regulatory elements and how they are associated with a phenotype. The RERs of a particular gene are measured by examining the rate of nucleotide substitutions over time on each branch of a phylogenetic tree. If the gene has fewer substitutions than the branch average for all genes, the gene’s evolution is decelerated on the branch, and the RER value is negative. If there are more substitutions than the average for the branch, the gene’s evolution is accelerated, and the RER value is positive. This information can inform us about selection occurring on the gene: a gene with fewer substitutions is likely under stronger selective constraint whereas a gene with more substitutions is under relaxed selective constraint or positive selection. RERconverge is an R package that tests for associations between RERs and phenotype values to identify genes with convergent changes in relative rates of molecular evolution on lineages associated with a phenotype of interest (Kowalczyk et al. 2019). RERconverge then ranks genes based on the significance of the association between their RERs and the convergent phenotype, allowing users to make predictions about the genes’ functions.

In a previous update, changes were implemented to enable analysis of categorical phenotypes (Redlich et al. 2024). As genomic data suitable for RERconverge studies are now available for hundreds and even thousands of species, it became important to write new efficient algorithms that could handle these large datasets. Also, as phenotypes become more diverse and complicated in such large phylogenies, it is important to provide finer control of the association tests to be performed. Here, we present updated methods with substantial increases to the speed of operation and improved statistical tests and outlier control.

## Materials and Methods

### Major Updates

The main workflow for RERconverge is as follows:

- *readTrees* takes a user-provided file with topology-constrained gene trees as input and reads the trees into a trees object
- *getAllResiduals* uses the trees object produced by *readTrees* to create a residual matrix of RERs for all branches present in all gene trees
- The user supplies phenotype values for extant species in the tree to generate a phenotype vector encoding this information for all branches (inferred using *char2Paths* or *tree2Paths*)
- *getAllCor* uses the phenotype vector and the residual matrix to test for a significant association between RERs and the phenotype for each gene

The primary runtime reduction for the *readTrees* and *getAllResiduals* functions resulted from an optimization of internal functions that iterate through data objects (these functions can be found in the *RERfuncs*.*R* source file, denoted with TT). *readTrees* was modified to contain the argument *useSpecies*, which allows users to filter their data to a specific subset of species, and more descriptive errors and warnings were added.

The function that calculates RER values, *getAllResiduals*, is now split into three sequential sub-functions to add finer control and better code readability. The first function, *transformPaths*, applies a transformation (square root by default) to the branch lengths of the gene trees, to reduce the effect of heteroskedasticity (Partha et al. 2019). The RER calculation function, named *coreGetResiduals*, takes the resulting modified trees object and calculates relative evolutionary rates for all branches of each tree as described in Partha et al. (2019). Finally, the function *getRMat* transforms the RERs output from the *coreGetResiduals* function into an RER matrix compatible with previous versions of RERconverge. The *weighted* argument was split into two arguments, *use*.*weights* and *weights*, for clarity and ease of use, but the default is still to calculate weights and perform weighted regression as in the previous version of RERconverge. The resulting data object is of the same type and size, so compatibility with previous versions is maximized.

The *getAllCor* function gained two outlier control options for weighted correlations to reduce the influence of extreme outliers. The first option, Winsorization, brings the *n* most extreme (highest and lowest) values to their *n*+1th most extremes. For example, when Winsorizing with a value of 3, the highest three values are set to the fourth highest, and the lowest three are set to the fourth lowest. Winsorization was previously only used in unweighted correlations but now is applicable to both weighted and unweighted. The other method for outlier control is bootstrapping, wherein the branches of a gene tree are resampled with replacement (1,000 iterations by default) and the association statistic is calculated for each sample. These statistics calculated using the resampled branches are then averaged, reducing the effect of any one outlier. Bootstrapping also corrects for heteroskedasticity. For outlier control, we recommend using a *winsorizeRER* and *winsorizeTrait* value of 3 (this is the default). Additional changes to *getAllCor* include improved user warnings and an internal reorganization for streamlining future additions.

### Minor Updates

#### Available statistical methods

The initial publication of RERconverge included two statistical tests for associating RERs with phenotypes, Pearson’s correlation and Kendall’s Tau, which work best for continuous data or binary data, respectively, with a balanced distribution of foreground clades. Here, foreground refers to species and branches possessing the trait of interest. We added additional statistical methods to test for associations between RERs and different types of trait data, thus expanding the datasets that can be analyzed with RERconverge (Table 1, Fig. 1). For continuous data, Pearson, Spearman, and Kendall’s Tau correlations are optimal. With categorical non-binary data, ANOVA and Kruskal-Wallis associations are optimal. For binary traits, Pearson and Kendall’s Tau are appropriate given balanced foreground data. However, if one foreground clade is larger than others, it will have a larger influence on the results. To counteract this, weights are an option for a Pearson correlation and can reduce the effect of imbalance. Weights are automatically computed by the function as the reciprocal of the number of species in a foreground clade, as this spreads influence evenly across independent clades. A decision tree to aid in selecting the most appropriate method for a dataset is provided in Fig. 1.

**Table 1:**
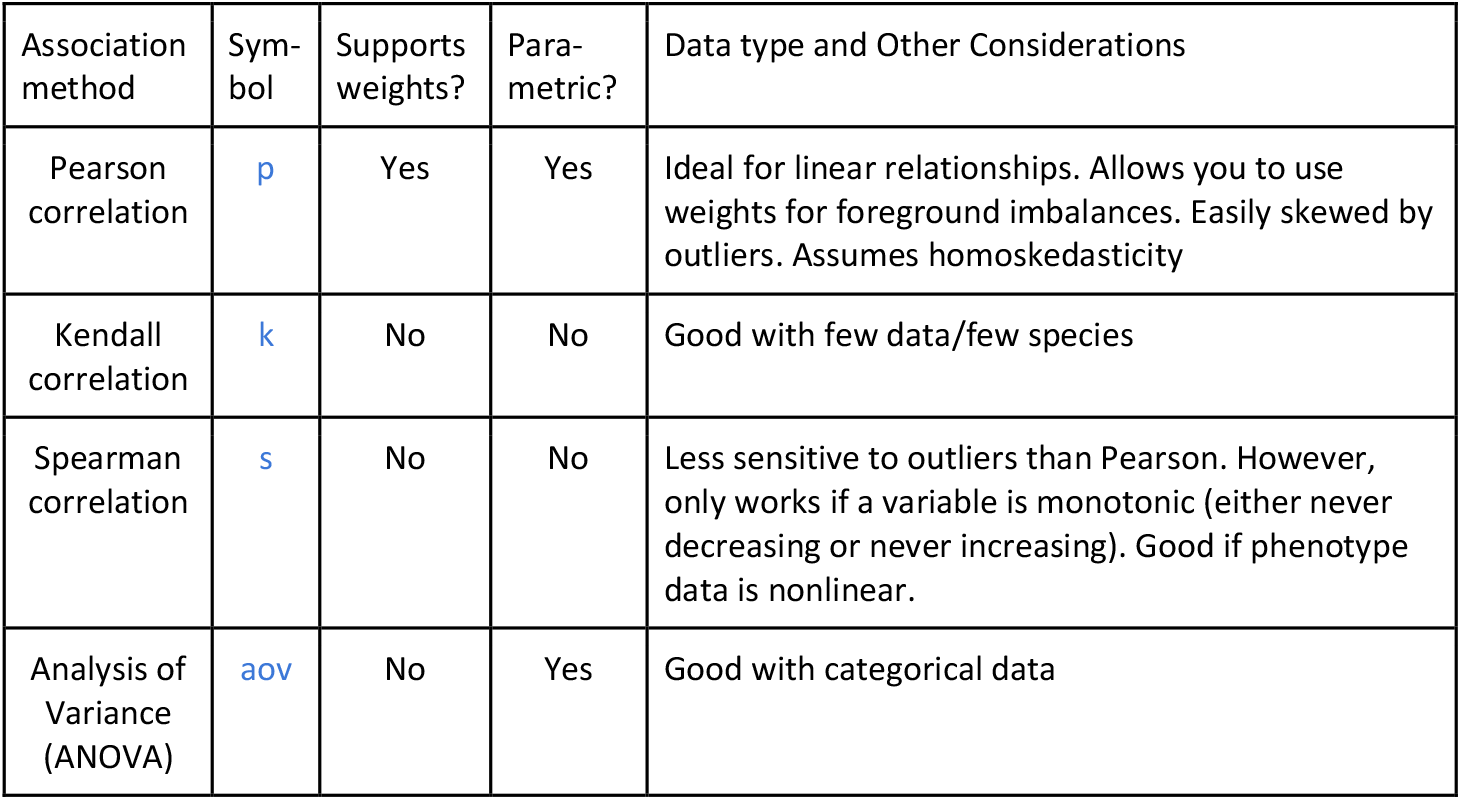
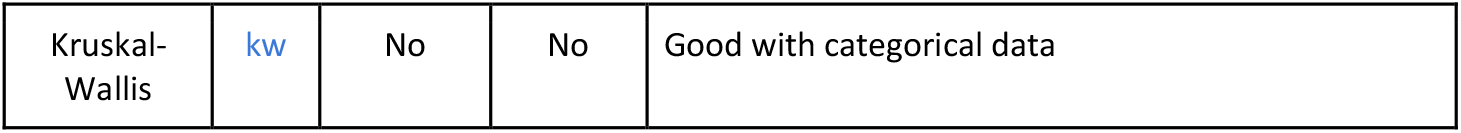
Statistical methods and their use cases.

| Association method | Sym-<br>bol | Supports<br>weights? | Para-<br>metric? | Data type and Other Considerations |
| --- | --- | --- | --- | --- |
| Pearson correlation | p | Yes | Yes | Ideal for linear relationships. Allows you to use weights for foreground imbalances. Easily skewed by outliers. Assumes homoskedasticity |
| Kendall correlation | k | No | No | Good with few data/few species |
| Spearman correlation | s | No | No | Less sensitive to outliers than Pearson. However, only works if a variable is monotonic (either never decreasing or never increasing). Good if phenotype data is nonlinear. |
| Analysis of Variance (ANOVA) | aov | No | Yes | Good with categorical data |
| Kruskal-Wallis | kw | No | No | Good with categorical data |

**Fig. 1:**
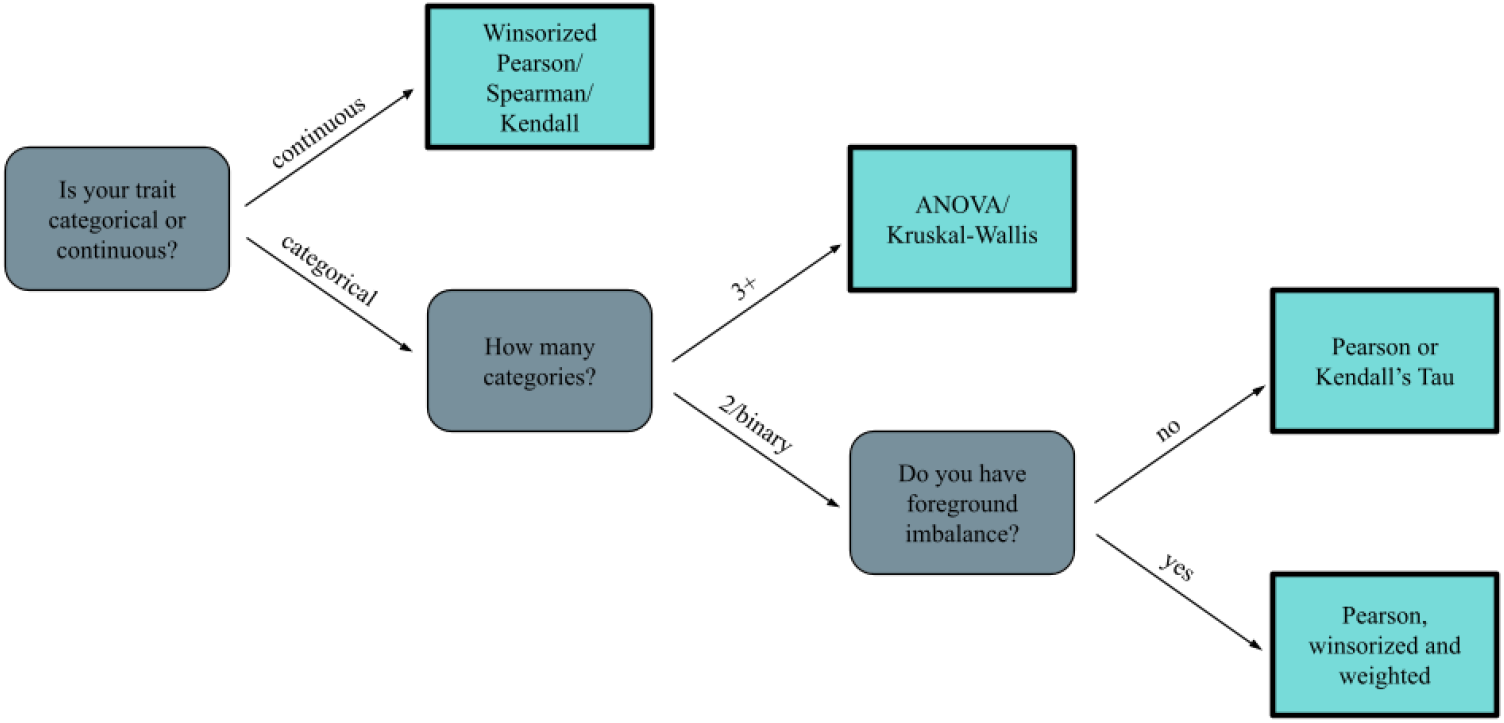
Decision tree for which statistical method to use with *getAllCor* based on phenotype data.

#### Changes to permulations

We previously described a method for testing the robustness of RERconverge results using a combined permutation- and simulation-based approach with data simulated over a phylogenetic tree (i.e., permulations (Saputra et al. 2021)). We introduced two versions of permulations for binary traits: (1) complete case (CC), which generates simulated data just once for every species present in the master tree, and (2) species subset match (SSM), which generates simulated data for each gene tree based on the subset of species present in the given gene tree. We have since introduced methods for running RERconverge with categorical traits (Redlich et al. 2024). Therefore, the CC permulations were updated to use the simulation used for categorical data, because these are compatible with both categorical and binary trait data. The new CC permulations maintain compatibility with existing input and output formatting. In the new version, setting *permmode* to “cc” will run this updated version of permulations, and the old version can be accessed using *permmode* “ccLegacy”.

The previous version of SSM permulations required simulated phenotypes to exactly match the foreground structure of the real data. For example, if the input data had exactly two pairings of sister species in the foreground, the simulated data needed to match this structure. For larger datasets and more complex tree structures, such as data where entire clades containing multiple species are in the foreground, the number of possible simulated foregrounds that match the topology of the real data is limited, and simulations rarely or never generated an acceptable foreground dataset. Therefore, we relaxed the requirement that simulated data must match the exact foreground topology of the real data. In the new version of SSM permulations, simulated trees must have the same number of extant foreground species (i.e., terminal branches) as the real data, but the evolutionary relationships among these foreground lineages are not restricted. Additionally, the previous version of SSM permulations often generated a biased distribution of foreground datasets, with some lineages almost always ending up in the foreground and others never appearing in the foreground in simulated datasets (Fig. 2A). This problem was particularly common when data were simulated over trees that had high variation in branch lengths, because phenotype values simulated over longer branches were always more extreme and therefore these long branches usually appeared in the foreground. To address this issue, the new SSM permulation function simulates data over a midpoint-rooted master tree, which results in a more random distribution of simulated phenotype values across the tips of the tree (Fig. 2B). The function subsets the master tree to only include species present in the gene tree but uses branch lengths from the master tree. In the new version of RERconverge, setting *permmode* to “ssm” will run the updated version of SSM permulations, and the old version can still be run by setting *permmode* to “ssmLegacy.”

**Fig. 2:**
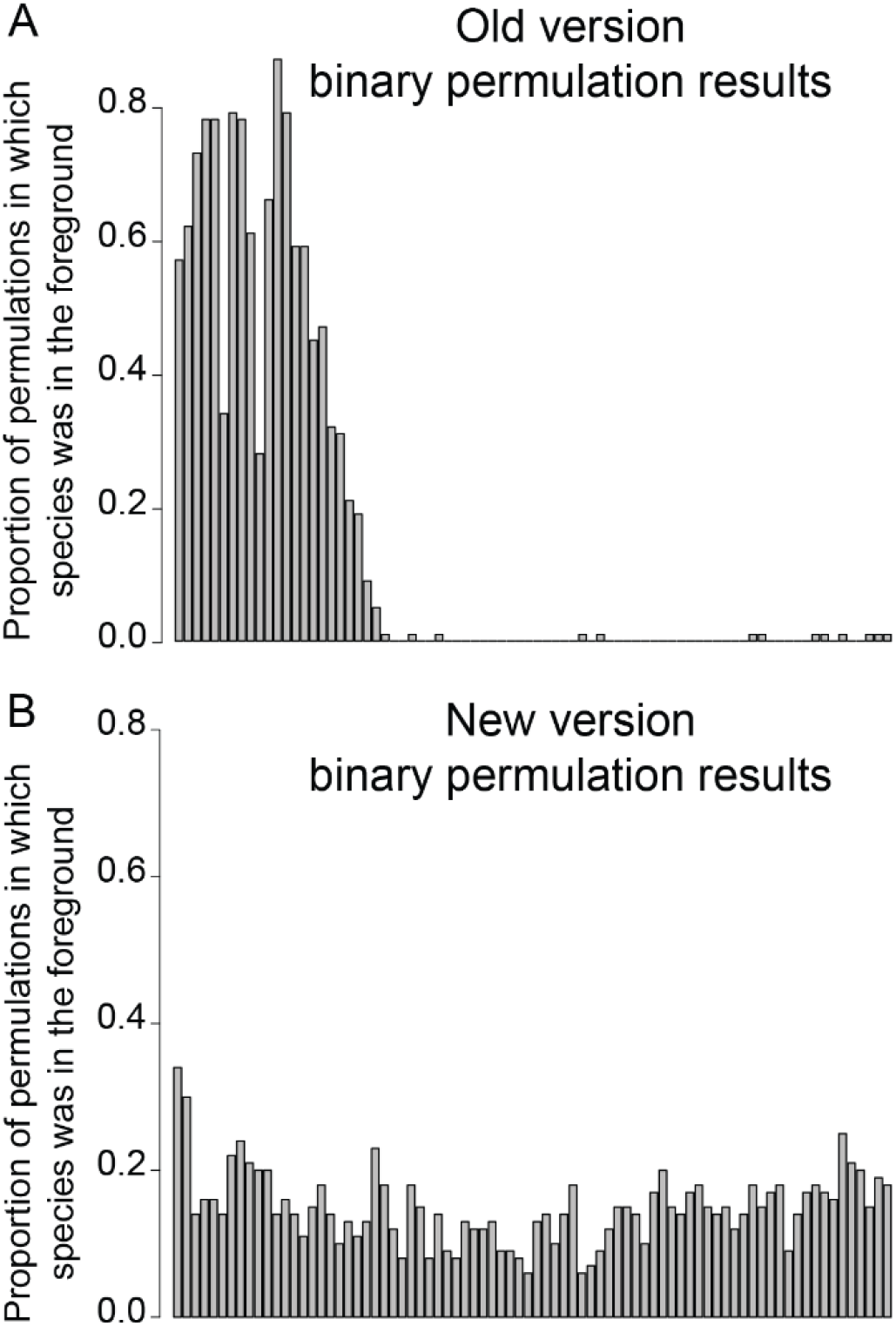
Updated SSM binary permulations show greatly reduced foreground bias. Plots show the proportion of permulations in which each species was assigned to the foreground based on simulated phenotype data using (A) the legacy version of SSM permulations (simBinPhenoSSM_legacy) versus (B) the updated version of SSM permulations (simBinPhenoSSM). Each bar represents a single species, and species names can be viewed in Supplemental Fig. S1. Each function was run using 100 permulations for a single gene (ENSMUSP00000001415; Data from (Kopania et al. 2025)).

#### Additional user-experience updates

Installation on Macs has been simplified due to the location of a universal GFortran library, meaning that binaries are no longer necessary; the installation instructions have been updated with the link to this library. Additionally, users can now modify species names in the plotting functions *plotRERs* and *plotTreesHighlightBranches*. Documentation and the user walk-through document have been updated accordingly. Lastly, many GitHub issues were solved over the course of development, generally yielding descriptive error messages to help users troubleshoot their data and results.

## Results and Discussion

### Substantially faster runtimes

The overhauled main workflow of RERconverge yields similar accuracy with a substantially faster runtime. The speedup factor represents how many times faster the new workflow operates in comparison to the old workflow and is calculated by dividing the old runtime by the new runtime. The improved efficiency was tested on both The University of Pittsburgh’s high performance computing cluster (HPC) and a personal computer (PC), with 80 trials for each combination of data file, pipeline, and system. Technical specifications of the systems are available in the Supplemental Information. On the HPC, the new workflow achieved sizable speedups starting at small amounts of data, but the speedup is especially notable as the number of gene trees increases. At 10,000 trees (with up to 62 species) using the HPC, on average, the new workflow takes 29 seconds, whereas the old workflow would take 563 seconds (9.5 minutes), a speedup factor of 19.1x (Fig. 3; Fig. 4). For datasets greater than 100 trees, the *readTrees* function reached speedups consistently over 50x on the HPC and 20x on the PC (Fig. 3). Speedup factors tended to increase as the number of input trees increased (Table 2). This drastic decrease in time for large datasets enables faster analysis and thus a reduced strain on computing resources, increasing accessibility of the package, and enabling analyses of massive collections of species genomes as are increasingly common.

**Table 2:** Times (in seconds) and speedup values for the high-performance computing (HPC) cluster and personal computer (PC) computation (mean of 80 trials).

| Tree counts | 100 | 500 | 1000 | 5000 | 10000 | 15000 | 19000 |
| --- | --- | --- | --- | --- | --- | --- | --- |
| HPC, original (s) | 7.14 | 40.15 | 69.72 | 279.01 | 562.56 | 987.45 | 1269.05 |
| HPC, new (s) | 1.67 | 5.57 | 9.70 | 21.78 | 29.37 | 39.65 | 44.32 |
| **HPC speedup factor** | **4.29x** | **7.20x** | **7.19x** | **12.81x** | **19.15x** | **24.90x** | **28.64x** |
| PC, original (s) | 5.79 | 25.82 | 4937 | 266.94 | 603.38 | 1010.97 | 1420.43 |
| PC, new (s) | 0.79 | 3.02 | 5.43 | 18.70 | 32.45 | 49.09 | 60.41 |
| **PC speedup factor** | **7.34x** | **8.54x** | **9.10x** | **14.27x** | **18.59x** | **20.59x** | **23.51x** |

**Fig. 3:**
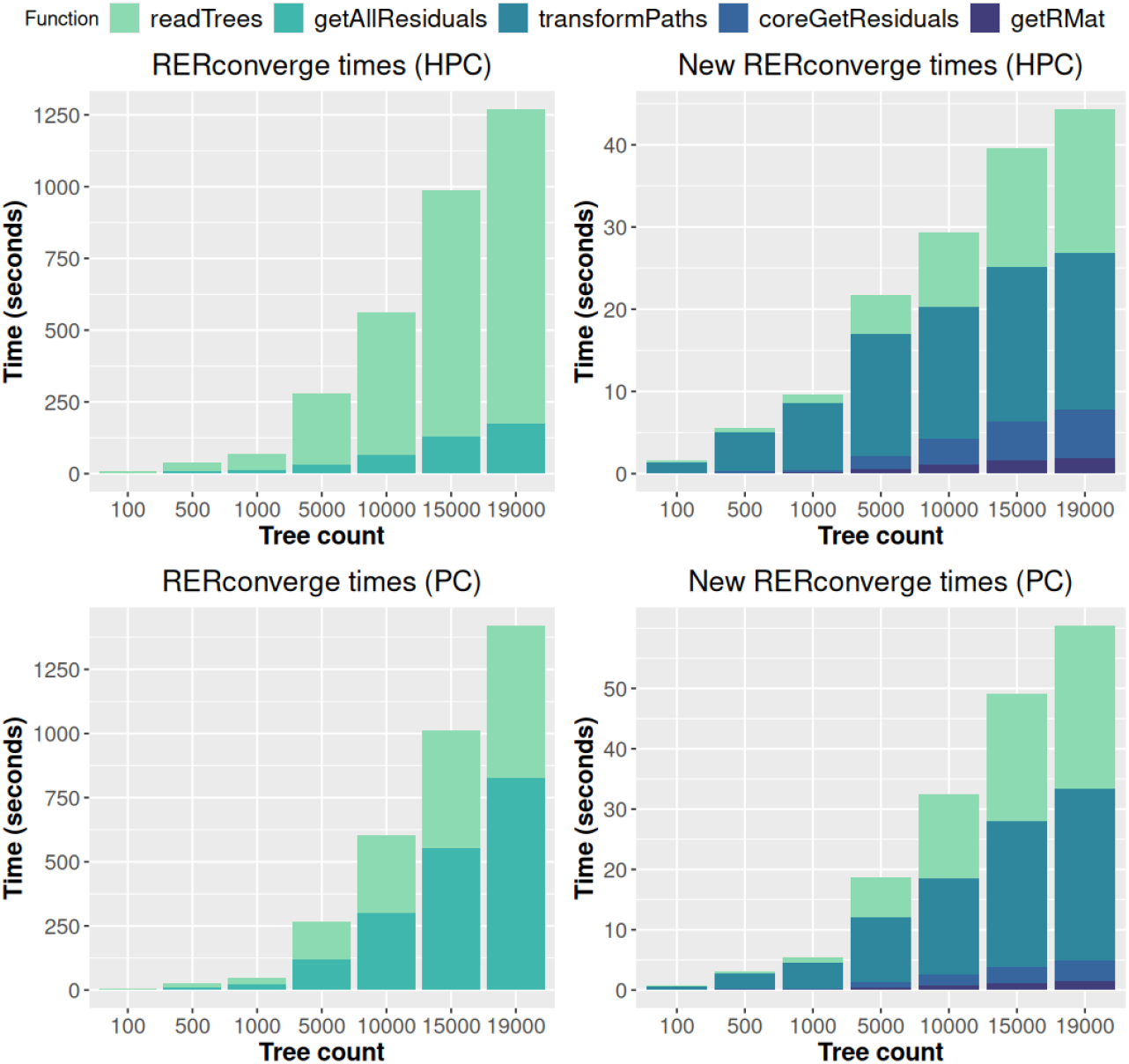
Runtime comparisons for the original and new RERconverge on a personal computer (PC) and the high-performance computing cluster (HPC). Note that the time axis labels differ between plots, which show a large decrease in total runtime for the new functions. The function bars show the proportion of runtime occupied by each function. Note that *transformPaths, coreGetResiduals*, and *getRMat*, take the place of *getAllResiduals* in the original runtime. Times are the means of 80 trials.

**Fig. 4:**
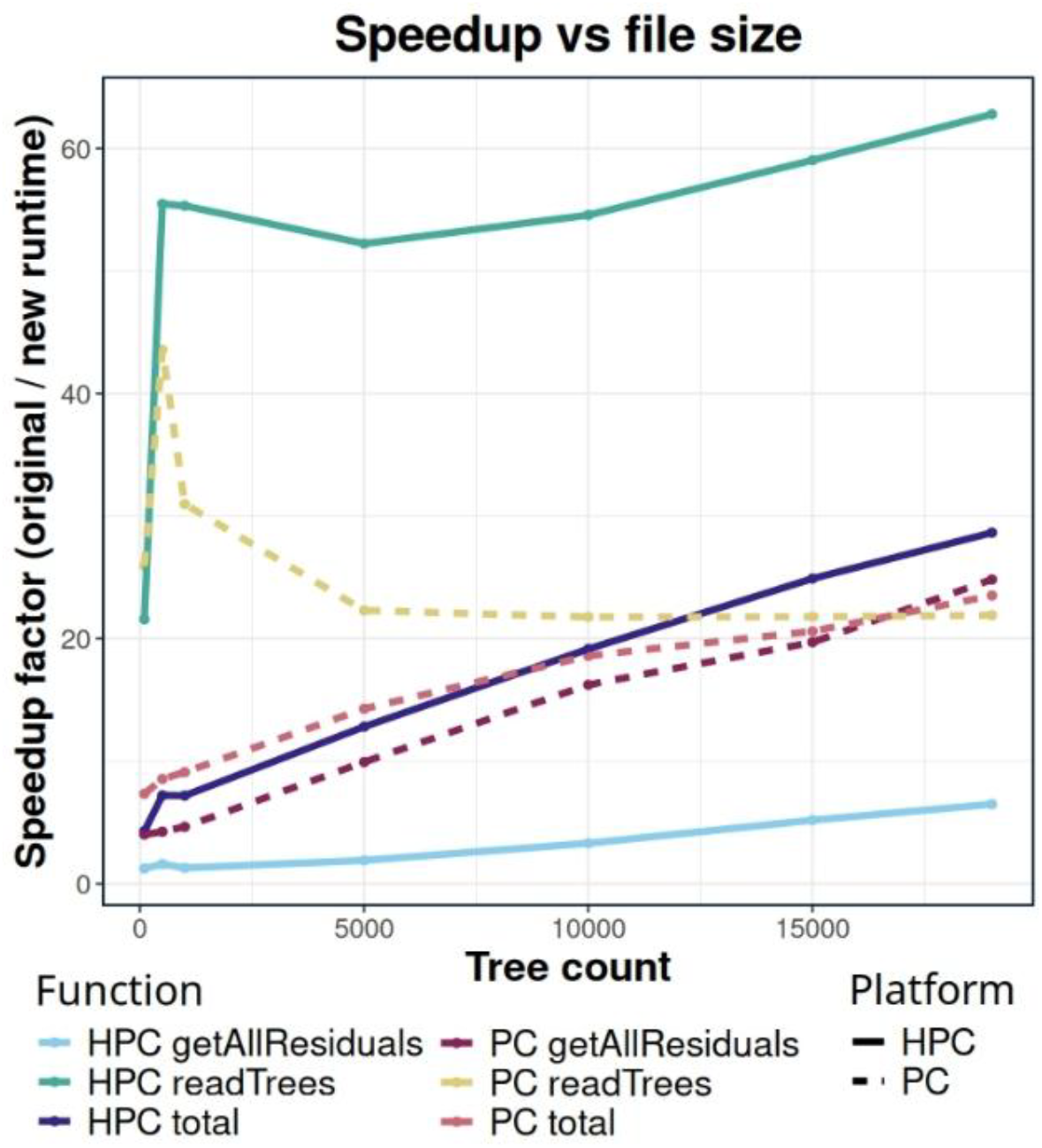
Speedup factor compared to the number of trees in each dataset. Separate speedups are presented for *readTrees* and *getAllResiduals*, as they make different contributions. Speedup factors represent the means of 80 trials.

### Consistency between old and new versions

#### readTrees

The bulk of the runtime reduction on the HPC was in the *readTrees* function (Fig. 3). To compare the original and new *readTrees* functions, we ran them on the same dataset and estimated the Pearson correlation between the output branch lengths from the two versions of the function. We used a dataset of 241 mammals (Genereux et al. 2020), and one of 343 yeasts (Shen et al. 2018). The resulting correlations had an R^2^ value of 0.999, indicating that the new *readTrees* does not sacrifice consistency for its improved runtime.

#### getAllResiduals

We tested the consistency of the new *getAllResiduals* algorithm with the old *getAllResiduals* algorithm using data from a study of convergent evolution in aquatic mammals, which is used as a tutorial dataset for RERconverge (Kowalczyk et al. 2019). Because the output from the new and old *readTrees* functions are nearly perfectly identical, we opted to use a tree object made using the new *readTrees* function. We ran this file into the old and new versions of the *getAllResiduals* functions, then ran the resulting residual matrices through *getAllCor* to correlate the outputs with phenotype values for aquatic mammals. We then sorted by the negative log of the p-value to generate two lists of genes ranked by acceleration. A Spearman correlation of these lists indicated a very strong correlation between the methods’ outputs (rho = 0.98, p < 1e10^−15^), indicating strong consistency between old and new results.

## Summary

RERconverge is a powerful tool to test for associations of gene evolutionary rate data with user selected traits of interest. By altering key internal functions, we achieved a much faster pipeline that produces results highly consistent with those generated by the previous method. Additionally, users have access to outlier control methods and more statistical methods than before to better handle different types and distributions of phenotype data. New permulation methods speed up runtimes with larger datasets and more complex tree structures, and these methods also generate less biased simulated data, even for trees with high variability in branch lengths. These improvements reduce the resources needed to request from computing clusters, reducing time and money spent and aiding users who have limited access to computing resources.

## Acknowledgements

We thank Matt Dean and members of the Clark Lab for feedback on changes as they were pushed to production.

