## Supplemental Information for "RERconverge Update: Runtime Reduction and Analysis Function Overhaul"

**Test specifications**

The improved efficiency was tested on both The University of Pittsburgh’s high performance computing cluster (HPC) and a personal computer (PC). We took a dataset of 19000 gene trees, and created seven data files with 100, 500, 1000, 5000, 10000, 15000, and 19000 trees. The HPC nodes are variable and have Intel Xeon Platinum 8352Y, Intel Xeon Gold 6248R, or AMD EPYC 9374F processors, and 8GB of ram per processor were allocated. The PC has an Intel i7 14700K processor and 32GB DDR5 RAM, running Linux Mint 22.1. Both HPC and PC used R 4.4, and 80 trials for each dataset and version were run.

**Supplemental Figures**

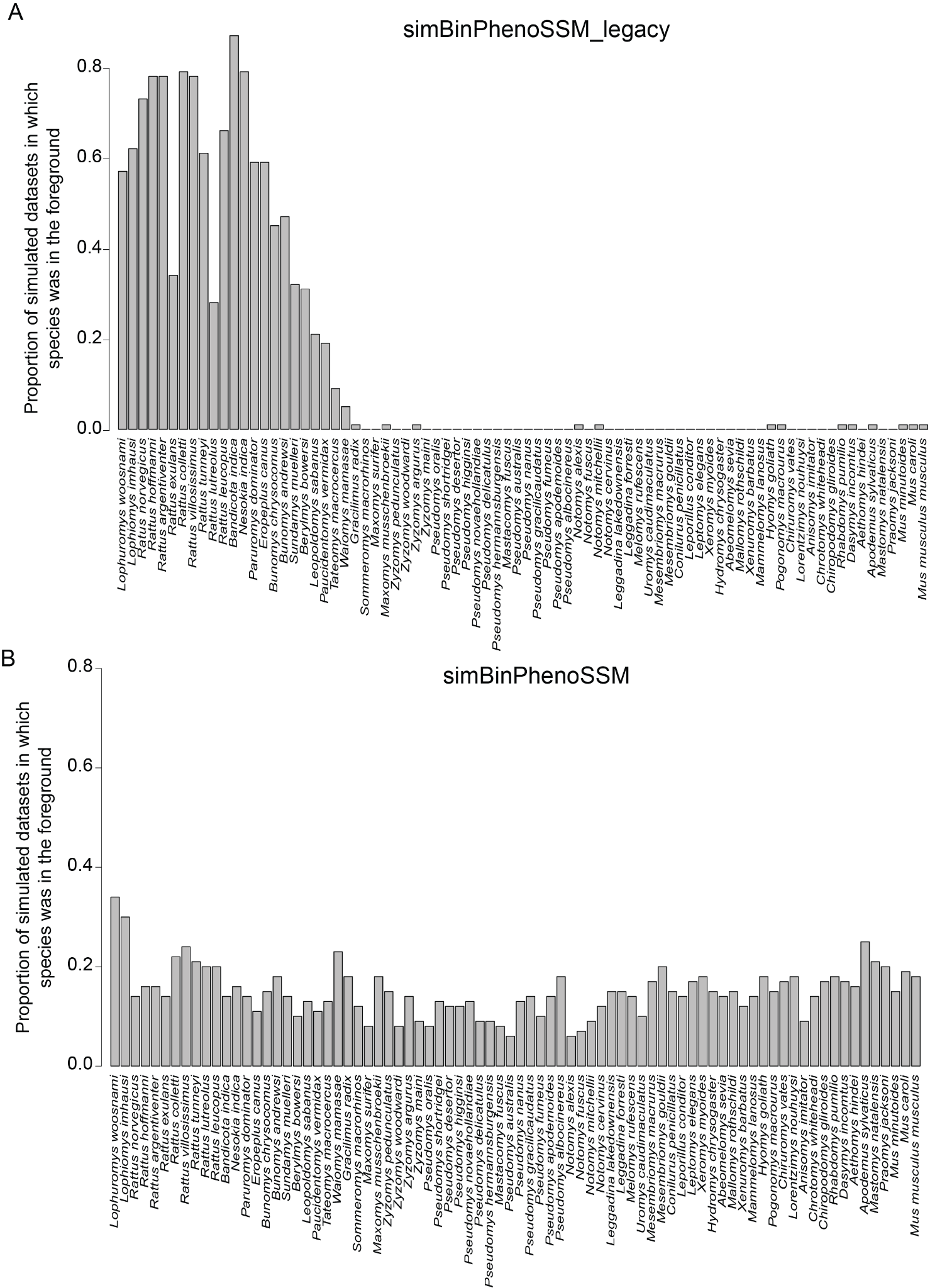

**Supplemental Figure S1: Updated SSM binary permulations show greatly reduced foreground bias.** Each plot shows the proportion of permulations in which each species was assigned to the foreground based on simulated phenotype data using (A) the legacy version of SSM permulations (simBinPhenoSSM_legacy) versus (B) the updated version of SSM permulations (simBinPhenoSSM). Each function was run using 100 permulations for a single gene (ENSMUSP00000001415; Data from (Kopania et al. 2025)).
